# Medium-assisted tumbling controls bacteria exploration in a complex fluid

**DOI:** 10.1101/2025.05.04.652133

**Authors:** Héctor Urra, Thierry Darnige, Xavier Benoit-Gonin, Justine Laurent, Angela Dawson, Wilson C.K. Poon, Eric Clément

**Affiliations:** Laboratoire PMMH-ESPCI Paris, PSL Research University, Sorbonne Université and Université Paris cité, 7, quai Saint-Bernard, Paris, France; SUPA and School of Physics and Astronomy, The University of Edinburgh, Edinburgh EH9 3FD, United Kingdom; Institut Universitaire de France (IUF)

## Abstract

In nature, many fluids that harbor bacterial populations or protect against microbial contamination exhibit non-Newtonian rheology. To study the spatial exploration of *E*.*coli* bacteria, a model multi-flagellated microorganism, in such complex environments, we design a motility medium with tunable macroscopic rheology. By increasing the solid charge in soft carbomer grains, we transition from a Newtonian viscous suspension to a yield-stress fluid. Using a 3D Lagrangian tracking device, we collect many individual bacterial tracks and characterize changes in motility properties such as swimming speed, persistence times and diffusivity for both a wild-type and a smooth runner mutant, up to the formation of a motility barrier at higher carbomer concentrations. We show that the presence of local mechanical disorder and resistance to penetration essentially override the biologically driven run-and-tumble navigation process. This “medium-assisted” exploration scenario is characterized by directional switching and stop-and-go kinematics and is closely related to the flexibility of the flagellar bundle.

## 1 Introduction

A crucial step in the bacterial life cycle is the planktonic state, where microbes spreading out from biofilm colonies autonomously explore their environment to eventually reach a new ecological niche ^1^. In many cases, the surrounding fluid exhibits non-Newtonian rheology, which drastically affects the exploration process. Moreover, to protect themselves from microbial aggression, many living species have developed physical defenses in the form of bio-gels, such as mucus, that cover the epithelial cells ^2–4^. In a different context, some soils or sandy clay layers are well known examples of natural filters capable of preventing microorganisms from contaminating freshwater reservoirs ^5–7^. In all of these examples, the issue of understanding the penetration of microorganisms in fluids with complex rheology is central. In a seminal paper, Wolfe and Berg ^8^ showed that the spreading of motile bacteria in semi-solid agar gel can actually be counterintuitive, as the best performing swarms were associated with an optimal tumbling frequency that allowed the cells to back away from obstacles in the agar matrix. From a fundamental hydrodynamic perspective, the question of swimming in non-Newtonian fluids has attracted considerable interest. For example, in viscoelastic media, propulsion via an undulating sheet ^9,10^ or via helical rotation ^11^ has been considered. Using molecular-based numerical simulations to model the surrounding polymer chains, the swimming performance of a model micro-swimmer was assessed ^12^. Many works suggest that there is a competition between elastic resistance to penetration and molecular organization around the swimmer, which ultimately increases propulsion efficiency. Experimentally, however, the situation is contradictory. Fluid elasticity has been found to either enhance ^13–18^ or hinder ^9,19–21^ swimming speed. Experimentally, swimming performance can depend on many details: bacterial shape, macroscopic fluid rheology, microscale mechanical disorder, and even the ability of the microorganism to directly metabolize small polymer molecules, which can induce a spurious increase in swimming speed ^20^. Interestingly, Patteson *et al*. ^18^ have studied a wild-type motile E.coli swimming in thin films of carboxy-methyl cellulose, a linear and flexible polymer. Increasing the polymer concentration induced faster and much more sustained movement, which was attributed to the suppression of the body’s wobbling rotation. Similar results were reported by Kamdar *et al*. ^22^ for *E*.*coli*, but here for bacteria swimming in a colloidal fluid. More recently, Bhattacharjee *et al*. ^23^ used a dense carbopol gel as a model complex environment to create an entrapped porous structure of microscopic hydrogel grains. They observed motile bacteria experiencing transient trapping and directional changes that characterize exploration of the highly confined pore space.

In this study, we investigate the motility of *E*.*coli* bacteria swimming in a model complex fluid with macroscopic flow properties ranging from Newtonian viscosity to yield stress Herschel-Bulkley rheology. To this end, we use a well-studied carbomer gel, i.e. a cross-linked polyacrylic acid polymer dispersed in a motility buffer at different concentrations ^24,25^. This simplified motility medium not only exhibits well controlled and tunable macroscopic rheology, but also introduces structural-mechanical heterogeneity at the microorganism level due to the arrangement of swollen hydrogel grains. At high carbomer concentrations, the medium undergoes a jamming process characterized by a rigid solid granular structure that restricts bacterial movement in the pore space, similar to the situation studied by Bhattacharjee *et al*. ^23^. However, at intermediate concentrations, the ability of the macroscopic swimmer to penetrate the fluid and eventually reorganize the hydrogel structures is likely to influence exploration performance. First, we follow the 3D spatial exploration of a ‘smooth runner’ strain in which the tumbling mechanism associated with the flagellar bundling/unbinding process is suppressed. In addition, using another smooth runner mutant, we were able to directly follow the dynamics of the body and the flagella. The motility characteristics are then compared with those of a wild-type strain that would undergo in a standard motility medium, a run and tumble kinematics. This study provides a comprehensive perspective on the emergence of a motility barrier for motile microorganisms in a model complex fluid environment.

## 2 Material and methods

### 2.1 Motile bacteria strains

Here we use motile *E*.*coli* strains, including both wild type (WT) and smooth runner (SR) mutants, all derived from the *E*.*coli* AB1157 WT strain. The primary strains of interest in this study are the wild-type RP 437 and its smooth runner mutant CR20, obtained by deletion of the CheY protein (ΔCheY). CheY is responsible for a change in the direction of motor rotation that triggers the tumbling process. To observe the flagella, we use a ΔCheY mutant, AD63, derived from the AD62 strain. In this case, the smooth-running mutant AD63 expresses E-GFP (enhanced green fluorescent protein) in the body and an Alexa-red conjugated ‘sticker’ protein on the flagella (see Junot et al. ^26^ for technical details). All bacteria were prepared by a standard procedure involving overnight growth in rich medium, collection at mid-growth, followed by washing and dispersion in minimal motility buffer. They were then added to the motility medium with controlled rheology. Detailed preparation protocols can be found in the Supplementary Information document.

### 2.2 Controlled swimming medium and observation cell preparations

The motility medium consists of a mass *M*_*C*_ of dry carbomer powder (commercial Carbopol 940 from Fischer Scientific) thoroughly mixed with a Berg motility buffer (BMB) of mass *M*_*B*_. The mixture results in a carbomer mass concentration (*C*), expressed as a percentage (%) and given by 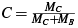. In all experiments, the swimming medium is kept at a pH of *pH* = 7 ± 0.2 and the temperature at *T* = 25^°^*C*.

For each experiment under the microscope, 5*µL* of dilute bacterial suspension in BMB (*OD* = 0.2) is gently mixed with 1*mL* of carbomer at a concentration *C* to obtain a highly diluted suspension that contains less than 3 × 10^7^ bacteria per mL.

A drop of 80*µ*L of the medium containing the motile bacteria is then placed in the experimental observation cell, which consists of two parallel glass slides enclosing a cylindrical drop of 1*cm* diameter with a confinement height of *h* = 230*µ* achieved by inserting a hollow double-sided tape between the slides (see Supplementary Information for details).

### 2.3 3D tracking and assessment of bacterial trajectories

The samples of carbopol mixtures containing the bacteria were observed using an inverted epifluorescence microscope (Zeiss Observer Z1) equipped with a C-apochromat objective (63 ×/1.2W). The individual 3D motion of fluorescent bacteria was monitored using an in-house Lagrangian tracking device, as previously described ^27^, performed using a Hamamatsu camera (ORCAFlash 4.0, C11440) at a frame rate of 80 FPS. This Lagrangian tracking technique has been used in previous studies to monitor *E*.*coli* swimming in different fluid environments, including quiescent fluid ^28^, Poiseuille flows ^29^, and surfaces ^26,30^ . For this “one-colour” tracking setup, we used the wild-type *E*.*coli* strain (RP437) and its Δ CheY smooth-runner mutant (CR20), both expressing a yellow fluorescent protein (YFP). For ‘two-colour’ tracking, the setup was extended using a beam splitter device to allow Lagrangian tracking of the bacterial body in green and simultaneous monitoring of the flagella in red.

## 3 Onset of a motily barrier

First, we collected trajectories of the “smooth swimmer” mutant CR20 in different carbomer mixtures, with concentrations ranging from C = 0 to C = 0.5% (four examples of such trajectories are shown in Figure 1A). Notably, the higher concentrations we use, corresponds to the starting point of the motility study proposed by Bhattacharjee *et al*.. ^23^ (note that their commercial carbopol product is slightly different). At each concentration, we obtain between 20 and 30 individual tracks at least 10 *µ*m away from the walls. In the standard motility buffer (*C* = 0), the trajectories are essentially persistent (see panel I of Fig.1A) with continuous changes in orientation due to Brownian orientation diffusion (diffusion coefficient *D*_*r*_ = 2(±0.8)10^−2^*s*^−1 28,32^). By post-processing the trajectories, we extract the time-resolved swimming velocities *V*_*b*_(*t*) (Fig.1B) and access the corresponding velocity distribution over the tracking times (Fig.1C). The mode of the velocity distribution (if available) gives a typical swimming velocity. Averaged over individual tracks, the mean swimming velocity in the motility buffer is 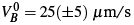 *µ*m/s. However, in the carbomer mixtures, the trajectories show frequent changes in direction as well as sequences of transient stops (Fig.1A and B). As the carbomer concentration is further increased, a large fraction of the bacteria exhibit significant wobbling but remain essentially trapped (see Fig.1A, panel IV) during the time window of our observations. When all encountered bacteria are trapped, the motility barrier is reached. The progressive changes in motility characteristics with concentration are associated with an evolution in instantaneous swimming speeds along the tracks (Fig.1C). It evolves as a progressive appearance of transient trapping periods characterized by low speed values, which eventually coexist with moments of ballistic motion. At higher concentrations, the ballistic motion disappears and is replaced by jumps in velocity. Eventually, a state is reached where all bacteria in the medium are trapped with almost no apparent motion. The medium then acts as a motility barrier. These changes in velocity fluctuations are also associated with a geometrical evolution of the track shape. Two parameters are used to quantify this evolution. The first is the compacity number, denoted as *N*_*c*_ = 2*R*/*L*, introduced by Miño *et al*. ^33^, this parameter is defined as the ratio of the maximum exploration diameter (2*R*) to the total track length (*L*). *R* is obtained as the radius of the minimum sphere encompassing a track. For linear tracks *N*_*c*_ → 1. For diffusive tracks *N*_*c*_ ∝ *L*^−0.5^ and for localized tracks *N*_*c*_ ∝ *L*^−1^, both tending to zero for long tracks.

**Fig. 1.**
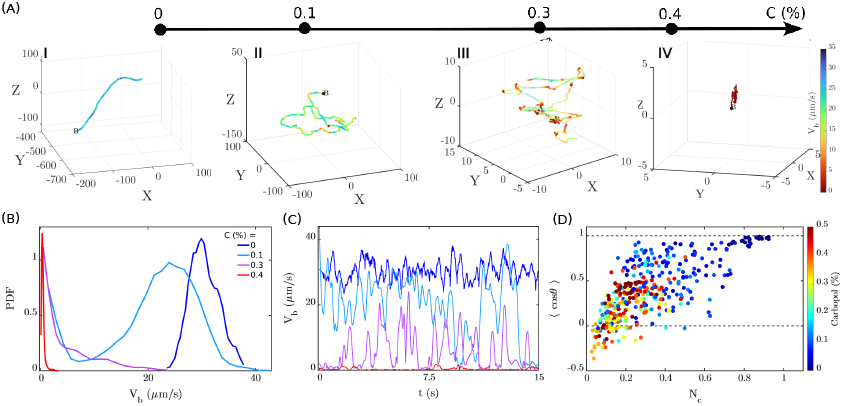
Onset of a motility barrier for non-tumbling *E*.*coli* mutants (CR20) exploring a carbomer mixture of increasing mass concentration *C*. (A) Examples of trajectories for *C* = 0 % (I), *C* = 0.1 % (II), *C* = 0.3 % (III) and *C* = 0.4 % (IV), scales in microns. Label *B* indicates the beginning of the track. Colors correspond to swimming velocities *V*_*b*_(*t*). (B) Velocity distributions along the tracks. (C) Corresponding velocity PDFs (rescaled for better visualization). (D) Characterization of the geometrical evolution of the tracks in the “compacity” (*N*_*c*_), “swimming persistency” (⟨*cosθ* ⟩) plane. Symbol colors correspond to *C* values.

The second parameter used the “swimming persistency”, is characterizing the swimming direction angular persistence during the exploration process ^34^. Successive reorientation angles *θ* (*t, δt*) are computed for angular changes in swimming direction between *t* and *t* + *δt*. We compute for *δt* = 0.2 s the mean value ⟨cos *θ* ⟩ averaged along the track.

In Fig.1E, each track is shown as a concentration-coded color symbol. We visualize a progressive transition from straight persistent tracks (*N*_*c*_ ≈ 1 and ⟨cos *θ* ⟩ ≈ 1) at low concentrations to compact random exploration at high concentrations (*N*_*c*_ ≈ 0 and ⟨cos *θ* ⟩ ≈ 0). At higher concentrations, the compacity number is also small, but for several tracks ⟨cos *θ* ⟩ was found to be negative, indicating compact localized trajectories with subsequent opposite swimming directions, corresponding to trapped bacteria rocking back and forth in a “cage”. It is noteworthy that, for the wild-type strain (RP437) which undergoes run and tumble exploration in the motility buffer, once Carbopol is added, the bacteria display motility characteristics similar to those of the “smooth-runner” mutant (see in SI, figure equivalent to Fig.1D, for the wild-type strain). Another crucial element that emerges from this characterization of trajectory shape is the importance of local mechanical disorder. Microscopic mechanical heterogeneity is manifested by a large variability in the values of *N*_*c*_ and < cos *θ* >, at a given concentration *C*. Note that, raw Carbopol products are commercially available as dry powders with a typical particle size of 0.2 *µ*m. In aqueous solvent the grains swell and the medium prepared by thorough mechanical mixing appears homogeneous and transparent. Here, at *pH* = 7, we could still identify weak optical heterogeneity as low contrast spherical patterns with a typical scale of 6 *µ*m (see SI), which corresponds to a typical bacterial size, including flagella.

## 4 Relating macroscopic rheology to bacteria exploration

### 4.0.1 Carbopol rheology

When Carbopol is added to the aqueous MB solvent, the mechanical properties in terms of flowability change drastically. In order to quantify at the macroscopic level these properties, we performed systematic rheological measurements to extract rheological parameters (see SI). We found for a Carbopol concentration *C* < *C*\* = 0.1% a Newtonian viscosity increasing with concentration (see Fig. 2A) as *η*(*C*) = *η*_0_(1 + *C*/*C*_1_ + (*C*/*C*_2_)^3^), with *η*0 = 1.06 · 10^−3^ Pa·s and *C*_1_ = 0.0337% and *C*_2_ = 0.0849%.

**Fig. 2.**
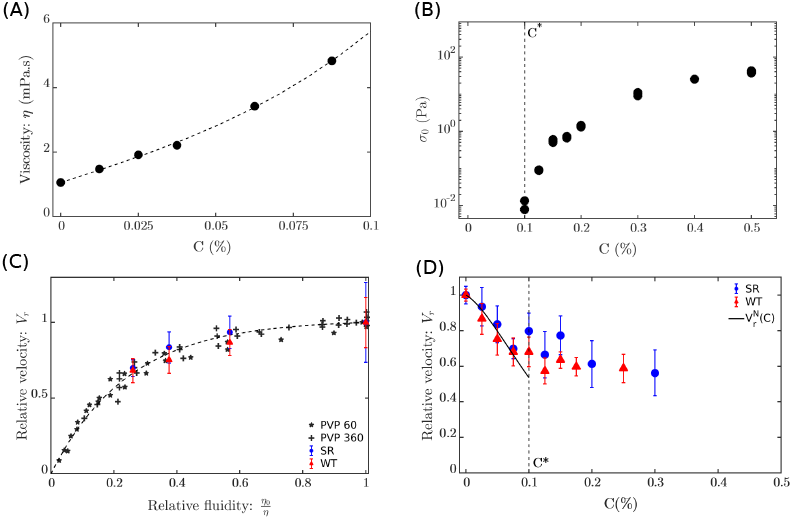
Influence of macroscopic rheological properties on the swimming speed of bacteria. A) Newtonian viscosity for Carbopol concentration *C* < ≤ *C* * = 0.1%, dashed line, third-order polynomial fit (see text). B) Yield stress values *σ*_0_(*C*). C) Relative swimming velocities *V*_*B*_/*V* ^0^ in the Newtonian regime as a function of relative fluidity *η*_0_/*η*. Light black symbols represent PVP60 and PVP360 solutions, data adapted from Martinez et al. 20. The dashed line is a one parameter fit of these data using eq.(2). Blue and red symbols are our data for smooth-runners (CR20) and wild-type swimmers (RP437).(D) Mean relative swimming speeds as a function of carbopol concentration: *V*_*r*_(*C*). The black line is the relationship *V*^*N*^ (*C*) (see e.g.3), the expected velocity reduction in a Newtonian fluid.

Above the concentration *C*\* we observed the onset of a yield stress. The rheology of Carbopol is then characterized by a nonlinear Herschel-Bulkey (HB) flow curve, well fitted by a relationship 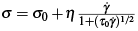(see SI). The corresponding yield stress increases with concentration as shown in Fig. 2B. Now we want to estimate the ability of a bacterium to penetrate such a yield stress fluid (i.e. when *C* > *C*^*^) by comparing the active pushing pressure *P*_*p*_ with the yield stress value. A pushing force *f*_*p*_ gives a stress *P*_*p*_ = *f*_*p*_/*a*^2^, where *a* is of the order of a bacterium’s body size. Its maximum value is related to the maximum active torque *T*_*M*_ exerted by the motor through the scaling relation *T*_*M*_ ∝ *f*_*p*_*a*. A dimensionless penetration number *N*_*p*_ can then be defined as the ratio of the shear stress to the macroscopic yield stress *σ*_0_(*C*):

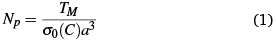

Interestingly, when applied to our E.coli strains, using a typical body size *a* = 2 *µ*m and a maximum motor torque *T*_*M*_ = 5 · 10^3^ pN·nm as reported in the literature (see Berg *et al*. ^35^), the crossover value *N*_*p*_ = 1 corresponds to a yield stress *σ*_0_(*C*_*M*_ ) ≈ 0.625 Pa, thus giving a carbomer concentration *C*_*M*_ ≈ 0.18 %, right in the motility transition region. Note however that Carbopol mixtures can be sensitive to aging. Our experiments are only marginally affected by this effect, as the samples were frequently replaced and the experiments did not last long enough for the rheological properties to vary more than 10% (see SI for aging test experiments on Carbopol blends).

#### 4.0.2 Swimming velocity

This change in rheological properties with concentration has a significant effect on the speed at which bacteria swim in the fluid. Martinez *et al*. ^20^ made systematic measurements of E. coli swimming in different polymeric fluids. They used polyvinylpyrrolidone polymer (C_6_H_9_NO)_n_, also known as PVP, with different molecular weights. At low concentrations, the polymeric fluids are essentially Newtonian, with viscosity increasing with concentration. Using a high throughput diffusion method (DDM) ^36^, they obtained the body rotation velocities as a function of polymer concentration. In Fig. 2C, we report their data replotted to show the relative swimming velocity as a function of the relative fluidity (defined as the inverse of the viscosity). To generate these data, we used the inherent linear relationship between body rotation rate and swimming speed at zero Reynolds number. The reference point for velocity 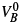 and viscosity 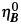 corresponds to the situation in the pure motility buffer. Black symbols on this graph correspond to PVP solutions tested by Martinez et al. ^20^ in the Newtonian regime with a viscosity increasing with the concentration. The relative velocity is fitted as a function of the relative fluidity: *x* = *η*_*O*_/*η* using an empirical function :

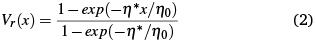

It is noteworthy that the velocity data for our bacteria (either smooth-runner or wild-type) quantitatively show the same dependence on relative fluidity and fall on to the velocity-fluidity master curve . We now have two empirical relationships for viscosity in the Newtonian regime as a function of the Carbopol concentration (See Fig. 2 A) and for relative swimming speeds as a function of fluid viscosity (eq.(2)). Thus, using both expressions, we derive an expression for the expected relative bacteria velocity as a function of the Carbopol concentration *C*:

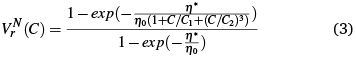

In Fig. 2 D shows the relative swimming velocity 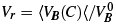 as a function of carbopol concentration for both the smooth and wild-type strains. For *C* < *C*^*^, i.e. in the Newtonian regime, the velocity data for both strains show a similar decrease, in quantitative agreement with the relationship (3) derived from Martinez *et al*. ^20^ on the expected decrease in swimming velocity with increasing viscosity (see black line in Fig.2D).

Above *C*^*^, the proportion of tracks where swimming speeds could actually be measured decreases. However, we did not observe a strong decrease in speed with concentration when it was possible to measure it. Rather, we observed bacteria swimming persistently at speeds comparable to those reached at the onset of the yield stress, but for shorter times.

#### 4.0.3 Swimming persistence time

For each trajectory, we extract a time-resolved swimming orientation along the trajectory 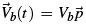, where 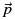 is the orientation unit vector. The orientation correlation function was obtained for a time lag 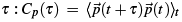, with a sliding average over time. Note that sometimes the dynamics can be very heterogeneous within the observation time window (stop and go dynamics). The maximum time lag value *τ*_*max*_ is chosen to be one tenth of the total track duration. For time lags in a window *τ*_*min*_ < *τ* < *τ*_*max*_, we fit *C*_*p*_(*τ*) to an exponentially decaying function to extract a typical swimming persistence time *τ*_*p*_. We extract *τ*_*p*_ values only when *τ*_*p*_ > *τ*_*min*_ = 0.5 s. In the absence of velocity persistence, we categorize the tracks are purely diffusive or eventually quasi-trapped. In Fig.3A we show the mean persistence times ⟨*τ*_*p*_(*C*)⟩ averaged over all tracks where it could be measured. It is known from previous work that in the absence of carbomer (*C* = 0)), the wild-type and smooth mutants have very different motility characteristics. The wild types show a wide distribution of persistence times resulting from behavioral variability ^26,28^. Smooth-runners experience a Brownian rotational diffusivity with a swimming persistence time of about 30s ^32^. Our measurements give a similar value (not shown in the graph). However, starting from the lowest concentration of carbomer used (*C* ≤ 0.025 %), a radical change in motility characteristics is observed for both strains. Fig.3A shows very similar mean persistence times for both the wild type and the mutant, which decrease monotonically with concentration down to the cut-off value of 0.5s. Note that even in the Newtonian regime (*C* < *C*^*^) all tracks show a persistence time above the cut-off value, whereas in the yield stress regime one observes the emergence of purely diffusive trajectories (*τ*_*p*_ < 0.5 s) as well as quasi-trapped trajectories. Above *C*\* one still observes many persistent trajectories up to concentration values slightly above 0.2%.

**Fig. 3.**
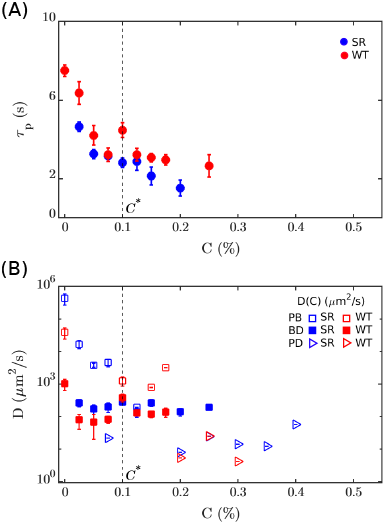
Characterization of the exploration process for *E*.*coli* wild-type (WT) and smooth runner (SR) strains. Blue and red colors correspond SR and WT respectively. (A) Mean persistence times ⟨*τ*_*p*_(*C*)⟩. (B) Mean spatial diffusivity ⟨*D*(*C*)⟩ for tracks identified in the three motility regimes PB (empty squares), BD (filled squares) and PD (triangles).

#### 4.0.4 Diffusivity

Finally, to characterize the exploration process upon reaching the motility barrier, we extract for each track (for SR and WT) the mean square displacement (MSD) *R*^2^(*τ*) by time averaging over the track duration. The limitations of the observation time do not allow us to draw conclusions about the actual long-term exploration process and the extent to which final capture may occur. However, these measurements reveal a qualitative evolution of the mesoscopic exploration process during the formation of the motility barrier. The MSDs were fitted with the two-parameter function: 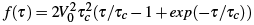, giving a diffusion coefficient 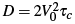. Only time lags *τ* smaller than a maximum value *τ*_*max*_, taken as 1/10 of the total track duration, were considered for the fit. At lower concentrations we encountered many tracks for which *τ*_*c*_ ≤ *τ*_*max*_, those in the observation range, are essentially ballistic and are termed “pure ballistic” (PB) (see Fig.1A, panel I, as an example). We associate with these tracks a proxy value for the diffusion coefficient 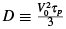 consistent with the standard picture of a random walk arising from run and tumble kinematics ^34^. We have found it to be a reasonable approximation when distinct ballistic and diffusive regimes are observed (see SI). These last tracks are called “ballistic diffusive” (BD) (see Fig.1A, panel II, as an example). Some tracks do not show any ballistic regime and the MSD is linear. These are called “purely diffusive” (PD) (see Fig.1A, panel III, as an example). Note that some tracks that show quasi-trapping during the observation time or a significant part of it would not fall into any of these categories (see Fig.1A, panel IV, as an example). On Fig.3B we show on a log scale for the three categories (PB, BD, PD) the mean diffusion coefficients ⟨*D*⟩ for different concentrations. Each category corresponds to well separated ranges of diffusivity values and the values obtained (10^3^*µm*/*s* for PB for *C* > *C*\*, 10^2^*µm*/*s* for BD and 10^1^*µm*/*s* for PD) do not seem to vary much with concentration. Moreover, for a wide range of concentrations, the three categories can coexist at the same concentration. All these features point to the influence of mechanical heterogeneity, which locally affects the exploration process. Furthermore, the quantitative correspondence between the diffusivity values of the smooth runners and those of the wild types indicates that the exploration process essentially ignores the internally driven tumbling process. Fig.3B also shows that (except for *PB* tracks), crossing the macroscopic stress onset at *C* = *C*\* does not lead to drastic changes in the diffusion coefficients. However, above a concentration of about ≈ 0.25%, only the purely diffusive tracks remain (as well as quasi-trapped bacteria). The diffusion values around 10*µm*^2^/*s* for *PD* tracks are significantly higher than the 2*µm*^2^/*s* maximum reported for the hoping and trapping process observed by Bhattacharjee *et al*. ^23^. The previous estimation of the shear stress seems to indicate that above this concentration the bacteria pushing pressure may not be large enough to reorganize the microscopic structures, however, as we will see in the next section, there may still be opportunities for bacterial movement, bridging the gap with the study by Bhattacharjee *et al*. ^23^, which essentially occurs at higher carbomer concentrations.

## 5 Medium assisted tumbling (MAT): an interplay between mechanical disorder and fluid-structure instability

The previous results have shown that the motility characteristics of wild-type and smooth runners become essentially the same when carbopol is added; the question now is to identify, from a micro-hydrodynamic perspective, what really controls the navigation process. Therefore, we turn to the “two-colors” tracking of the smooth runner mutant (AD63). Fig.4 shows two characteristic events leading to disorientation and trapping at concentrations *C* = 0.125 % and *C* = 0.25 %. In Fig.4A, a direct visualization of the body and the flagellar bundle for a smooth swimmer in parallel to the corresponding swimming trajectory, brings evidence for a change in swimming direction resulting from a deflection of the flagellar bundle with respect to the body orientation. In Fig.4B we show that a strong buckling process of the flagellar bundle leads to a drastic redirection of swimming. The corresponding movies can be viewed in SI. In the supplementary videos, we also show a situation at *C* = 0.25 % where a bacterium remains quasi-trapped for more than 7*s* and undergoes a release process activated by a sudden buckling of the flagellar axis with respect to the body direction. In fact, the connection between the flagellum and the body is mediated by a soft shaft called the “hook” ^37^. Recent numerical simulations of multiflagellated bacteria have explicitly shown how hook and flagellar elasticity are crucial for optimizing the bundling/unbundling process ^38^. Previous works have also proposed that hook and flagellar flexibility are crucial for inducing bacterial reorientation under shear ^39,40^ or for triggering tumbling of marine monoflagellated bacteria via a buckling process ^41^. More recently, frustrated tumbling has been shown to occur in liquid crystals ^42^ and to drive reversal motions of *E*.*coli*. All these examples highlight the central importance of the fluid-structure interplay between the elasticity of the flagellar apparatus and the mechanical properties of the surrounding fluid. In the present situation, we define “medium-assisted tumbling” (MAT) as the ability to navigate in a complex fluid and to overcome the intrinsic biological navigation mechanism via a fluid-structure interplay as described in Fig.4.

**Fig. 4.**
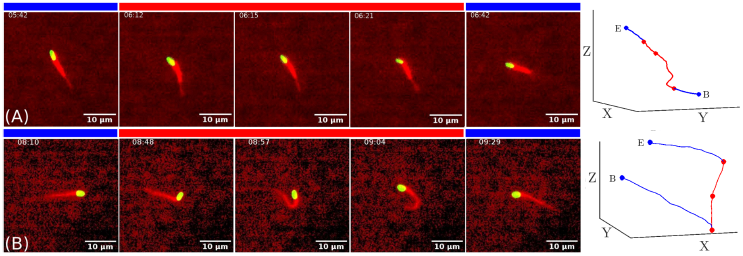
Visualization of flagellar bundle dynamics during “medium assisted tumbling” (MAT) using the “two-color” 3D tracking technique (Smooth Runner AD63 mutant). Left: time-lapse movie showing MAT dynamics. Right: corresponding 3D trajectories with time-lapse moments shown as dots. Label (B) (or (E)) represents the beginning (or end) of the trajectory. Blue bar and trajectory sections represent straight swimming directions before and after MAT. Red bar and trajectory sections represent changes in swimming direction due to MAT. (A): *C* = 0.125 % and (*B*): *C* = 0.25 %.

## 6 Conclusion

Using a model complex fluid with tunable macroscopic rheology, we monitored the formation of a motility barrier for motile *E*.*coli* bacteria exploring their environment. The model complex fluid is a motility buffer containing a solid charge of soft and swollen carbomer particles of typical bacterial size. Increasing the carbomer concentration allows the transition from a Newtonian fluid to a complex fluid exhibiting yield stress and Herschel-Bulkley rheology. As the macroscopic yield stress increases with concentration, an order of magnitude calculation of the resistance to bacterial movement shows that the ability to penetrate the fluid is inherently limited by the maximum internal torque that the bacteria motor can exert on the flagella. Therefore, at high concentrations of Carbomer, autonomous, self-propelled motility will inevitably cease and a motility barrier will be reached. However, from a microscopic perspective, the transition to complete arrest is a much more complex scenario. Using a 3D tracking technique, we observe and quantify the progressive evolution of the bacterial exploration kinematics and show how the addition of carbomer drastically affects the shape of the trajectories, promoting transient trapping episodes up to permanent trapping. We monitor the decrease in swimming speed, swimming persistence times and diffusivity for both a wild type and its corresponding smooth swimmer mutant (Δ*CheY* ). A remarkable result is that all measured properties are quite identical for the wild type and the smooth swimmer, suggesting that a media-related effect takes over the internal tumbling process. The presence, at a given concentration, of significant heterogeneity in the tracks, even before the macroscopic yield stress is reached, shows that the local mechanical disorder, probably resulting from the mesoscopic spatial structuring of the carbopol, is affecting the swimmer’s directional changes. Furthermore, direct visualization of the flagellar bundle during swimming, which shows bending (or even buckling) associated with changes in swimming direction, suggests an inherent interplay between the flexibility of the flagellar apparatus and the local mechanical landscape. This complex navigational process, which we term “Medium Assisted Tumbling” (MAT), is fundamentally different from the standard biological tumbling process. MAT induces a decrease in diffusivity of more than three orders of magnitude by reducing the persistence time, but also by controlling the trapping time. We have visualized that buckling-like changes in flagellar bundle shape are able to promote a trapping release process. This occurs at the higher carbopol concentrations and may be key to understanding the corresponding diffusion process consisting of transient hopping as described by Bhattacharjee et al. and visualized for carbopol concentrations corresponding to the higher limits of our study. This MAT scenario needs to be deepened and extended to other types of complex media and more generally, this process may become crucial in the context of chemotaxis ^43,44^, as the competition between internally driven vs. externally driven reorientation is key to understanding the efficiency of the chemotactic drift in confined and/or rheologically complex environments. Interestingly, in our situation, the active exploration of such a complex fluid affects the spatial diffusivity and swimming residence times in a manner opposite to the reports of Patteson *et al*. ^18^ for a bulk linear polymer matrix and Kamdar *et al*. ^22^ for a colloidal suspension. Therefore, further exploration of the MAT scenario to unravel the fundamental fluid-structure dynamics of soft and motile microorganisms in complex fluids could shed new light on the inherent complexity of the mechanical processes involved in microbial contamination.

## Author contributions

H.U. and E.C. designed the experiment . H.U. did the experiments and analyzed the data. T.D. did the 3D tracking algorithm and the numerical interfaces of the experimental set-up . X.B-G and J.L. contributed to the set-up construction and protocols for preparing Carbopol suspensions. A.D. did the E.coli AD mutants. H.U., W.P., E.C. discussed the scientific aspects and wrote the article.

## Data availability

Data for this article, videos and data sets are available at 10.5281/zenodo.15295338.

## Acknowledgment

We acknowledge the ANR PushPull (ANR-22-CE30-0038) grant and ANID BECAS CHILE 72180269 for H.U. Ph.D. grant. H.U. thanks H. Auradou and C. Manquest for help with the low shear rheometer. E.C and W.P. thank the CNRS/Royal Society Grant PHC-1576 and C. Douarche for providing *E*.*coli* RP437 and CR20 mutant. E.C. and H.U. thank T. Divoux for enlightening discussions on Carbopol rheology.

## Notes and references

1 R. Vasudevan, J Microbiol Exp., 2005, 1, 84–98.

2 M. E. Johansson, J. M. H. Larsson and G. C. Hansson, Proceedings of the national academy of sciences, 2011, 108, 4659–4665.

3 B. O. Schroeder, Gastroenterology Report, 2019, 7, 3–12.

4 N. Figueroa-Morales, L. Dominguez-Rubio, T. L. Ott and I. S. Aranson, Scientific reports, 2019, 9, 1–10.

5 G. Jiang, M. J. Noonan, G. D. Buchan and N. Smith, Australian Journal of Soil Research,, 2005, 43, 695–703.

6 R. R. E. Artz, J. Townend, K. Brown, W. Towers and K. Killham, Environmental Microbiology, 2005, 7, 241–248.

7 G. Fongaro, M. C. García-González, M. Hernández, A. Kunz, C. R. M. Barardi and D. Rodríguez-Lázaro, Frontiers in Microbiology, 2017, 8, year.

8 A. J. Wolfe and H. C. Berg, Proceedings of the National Academy of Sciences, 1989, 86, 6973–6977.

9 E. Lauga, Physics of Fluids, 2007, 19, 083104.

10 A. Kamal and E. E. Keaveny, Journal of the Royal Society Interface, 2018, 15, 20180592.

11 S. Gómez, F. A. Godínez, E. Lauga and R. Zenit, Journal of Fluid Mechanics, 2017, 812, year.

12 A. Zöttl and J. M. Yeomans, Nature Physics, 2019, 15, 554–558.

13 J. Teran, L. Fauci and M. Shelley, Physical review letters, 2010, 104, 038101.

14 B. Liu, T. R. Powers and K. S. Breuer, Proceedings of the National Academy of Sciences, 2011, 108, 19516–19520.

15 J. Espinosa-Garcia, E. Lauga and R. Zenit, Physics of Fluids, 2013, 25, 031701.

16 S. E. Spagnolie, B. Liu and T. R. Powers, Physical review letters, 2013, 111, 068101.

17 B. Thomases and R. D. Guy, Physical review letters, 2014, 113, 098102.

18 A. Patteson, A. Gopinath, M. Goulian and P. Arratia, Scientific reports, 2015, 5, 1–11.

19 X. Shen and P. E. Arratia, Physical review letters, 2011, 106, 208101.

20 V. A. Martinez, J. Schwarz-Linek, M. Reufer, L. G. Wilson, N. Morozov and W. C. K. Poon, Proceedings of the National Academy of Sciences, 2014, 111, 17771–17776.

21 B. Qin, A. Gopinath, J. Yang, J. P. Gollub and P. E. Arratia, Scientific reports, 2015, 5, 1–7.

22 S. Kamdar, S. Shin, P. Leishangthem, L. F. Francis, X. Xu and X. Cheng, Nature, 2022, 603, 819.

23 T. Bhattacharjee and S. Datta, Nature communications, 2019, 10, 2075.

24 D. Bonn, M. M. Denn, L. Berthier, T. Divoux and S. Manneville, Reviews of Modern Physics, 2017, 89, 035005.

25 M. Dinkgreve, M. Fazilati, M. Denn and D. Bonn, Journal of Rheology, 2018, 62, 773–780.

26 G. Junot, T. Darnige, A. Lindner, V. A. Martinez, J. Arlt, A. Dawson, W. C. K. Poon, H. Auradou and E. Clément, Phys. Rev. Lett., 2022, 128, 248101.

27 T. Darnige, N. Figueroa-Morales, P. Bohec, A. Lindner and E. Clément, Review of Scientific Instruments, 2017, 88, 055106.

28 N. Figueroa-Morales, R. Soto, G. Junot, T. Darnige, C. Douarche, V. A. Martinez, A. Lindner and É. Clément, Physical Review X, 2020, 10, 021004.

29 G. Junot, N. Figueroa-Morales, T. Darnige, A. Lindner, R. Soto, H. Auradou and E. Clément, EPL (Europhysics Letters), 2019, 126, 44003.

30 A. J. Mathijssen, N. Figueroa-Morales, G. Junot, É. Clément, A. Lindner and A. Zöttl, Nature communications, 2019, 10, 1–12.

31 R. e. a. Baillou, apreprint, ArXiv:2503.11364 [physics.bio-ph], 2025.

32 G. Junot, E. Clément, H. Auradou and R. García-García, Physical Review E, 2021, 103, 032608.

33 G. Mino, T. E. Mallouk, T. Darnige, M. Hoyos, J. Dauchet, J. Dunstan, R. Soto, Y. Wang, A. Rousselet and E. Clement, Physical review letters, 2011, 106, 048102.

34 P. S. Lovely and F. Dahlquist, Journal of theoretical biology, 1975, 50, 477–496.

35 H. C. Berg and L. Turner, Biophysical journal, 1993, 65, 2201–2216.

36 M. Reufer, V. A. Martinez, P. Schurtenberger and W. C. Poon, Langmuir, 2012, 28, 4618–4624.

37 H. C. Berg, E. coli in motion, Springer, New York, 2004.

38 B. Wu-Zhang, P. Zhang, R. Baillou, A. Lindner, E. Clement, G. Gompper and D. Fedosov, Journal of the Royal Society Interface, 2025, XX, xxxx.

39 M. Tournus, A. Kirshtein, L. Berlyand and I. S. Aranson, Journal of the Royal Society Interface, 2015, 12, 20140904.

40 M. Potomkin, M. Tournus, L. V. Berlyand and I. Aranson, Journal of The Royal Society Interface, 2017, 14, 20161031.

41 D. Brumley, R. Rusconi, K. Son and R. Stocker, The European Physical Journal Special Topics, 2015, 224, 3119–3140.

42 M. Goral, E. Clement, T. Darnige, T. Lopez-Leon and A. Lindner, Interface Focus, 2022, 12, 20220039.

43 D. Amchin, J. Ott, T. Bhattacharjee and D. Ss, PLoS Comput Biol, 2022, 18, e1010063.

44 M. Grognot, N. Woo, L. E. Elson and K. M. Taute, Proceedings of the National Academy of Sciences, 2023, 120, e2301873120.

