## Supplementary information for "Medium-assisted tumbling controls bacteria exploration in a complex fluid"

### PREPARATION PROTOCOLS

#### Preparation protocol for wild-type strains RP437 and smooth swimmer mutant strain CR20 (ΔCheY) in the motility buffer (body visualization only)

Bacteria are inoculated in 5mL of culture medium (M9G: 11.3 g/L M9 salt, 4 g/L glucose, 1 g/L casamino acids, 0.1mM $CaCl_2$, 2mM $MgSO_4$) with antibiotics (chloramphenicol at 25 $\mu$g/mL for RP437 an ampicillin at 100 $\mu$g/mL for CR20) and grown over night at 30 °C until early stationary phase. The growth medium is then removed by centrifuging the culture and removing the supernatant. The bacteria are re-suspended in a Motility Buffer (MB: 0.1mM EDTA, 0.001mM l-methionine, 10 mM sodium lactate, 6.2 mM $K_2HPO_4$, 3.9 mM $KH_2PO_4$) with 0.005 % of polyvinyl pyrrolidone (PVP) and is supplemented with 0.08 g/mL of L-serine. The addition of L-serine increases the bacteria mobility and PVP is used to prevent bacteria from sticking to the surfaces.

#### Preparation protocol for smooth swimmer mutant strain AD63 (ΔCheY) in the motility buffer

Bacteria AD63 stems from the AD62 strains developed in the group of Pr Wilson Poon in Edinburgh by Dr Angela Dawson. The preparation protocol is described in [1]. AD63 is the smooth-runner ΔCheY mutant of this strain. Both strains were constructed out of E.Coli AB1157 [2]. The wild type fliC gene, which encodes the flagellin sub-unit (which polymerises to form the bacterial flagella filament) was modified so that a cysteine was substituted for a serine amino acid at position 219 (S219C). This mutation allows a labeling of the flagella with Alexa Fluor 647 dye. Suspensions of AD63 are prepared using the following protocol: bacteria are inoculated in 10mL of Lurial Broth (LB) with ampicillin at 100 $\mu$g/mL and grown over night at 30 °C. Then 100 $\mu$L of this solution is inoculated in 10mL of Triptone Broth (TB) and grown during several hours until early stationary phase. The growth medium is then removed by centrifuging the culture and removing the supernatant. The bacteria are re-suspended in 1mL of Berg Motility Buffer (BMB: 6.2 mM $K_2HPO_4$, 3.8 mM $K_2PO_4$, 67 mM NaCl, and 0.1 mM EDTA) with 10 $\mu$L of Alexa red colorant (Alexa 647 at 5 mg/mL diluted in DMSO) and let under gentle shacking during 2 hours. The solution is then washed by centrifuging the culture and removing the supernatant part.

#### Carbomer mixtures

The carbomer used is Carbopol 940. The mixture preparation consists in dissolving the powder in the motility buffer (MB) via an helical mixer rotating for long times within a closed box to prevent drying. We initially prepared a mother solution at concentration $C = 0.5$ % and then dilute this sample to obtain the desired concentration values. For 50 ml of a Carbopol mixture, we use the following recipee:

- In a 150 ml falcon tube, we pour 50 ml of MB (Mobility Buffer) and slowly drop a mass $m_C = 0.25$ g of Carbopol powder in order to achieve the required mass fraction $C = 0.5$ %. While pouring, we mix in the Falcon bottle with the helical mixer at 60 rpm. The dispersion is done with a spoon pouring small amounts of Carbopol powder at a time, and waiting for its dissolution in the solution. This process is very slow and should be done over a period of half an hour.

- The solution is mixed for two hours at 60 rpm, under cover, until all the clumps are completely dissolved in the MB solution. At the end, one should obtain a semitransparent viscous substance of gray color.

- Thereafter, 150$\mu$/L of Triethanolamine (TEA) at concentration $\geq 99$ % is added into the solution and mixed to homogenize the gel. TEA is a biocompatible weak base that also helps clarifying the solution and does not change the pH (Standard TEA buffer is at pH $= 7.5$).

- At the end of the process, one should verify that the solution stays at pH$= 7 \pm 0.2$.
- Subsequently, we leave the mixer in rotation under cover for at least 48 hrs at 60 rpm. After this time, the pH should be stable at the desired pH$= 7 \pm 0.2$ value (control is done after the process).
- For the final dilution process to achieve a concentration $C$, the solution is mixed with a vortex mixer for 120 s.
- The final step consists in centrifuging the sample at 1350 rpm during 120 s, in order to remove any micro bubbles eventually created.

Hence prepared, the mother solution and the mixtures can be used over few day if stored in a closed container at a temperature of 4°C. When conducting the experiments, the sample must stands at room temperature to reach the working temperature of 25 °C.

### MACROSCOPIC RHEOLOGY

A low-shear rheometer (Contraves 30) with a cylindrical Couette geometry (external radius $R_i = 55$ mm and gap space $e = .5$ mm) and a standard rheometer (Anton Paar Physica MCR 501) with a cone-plate geometry (diameter of 25 mm) and. The experiments are performed at a temperature $T = 25$ °C. Carbomer mixtures put in the rheometer were initially sheared at high shear rate during 1 min (60 $s^{-1}$ for the low shear apparatus and 100 $s^{-1}$ for the cone plane geometry).

#### Low-shear rheometry

The range of concentrations used for the Carbopol mixtures starts from dilute solutions with a viscosity near water values. The low-shear Couette rheometer is then used to obtain the flow curves $\sigma(\dot\gamma)$ in a shear rates range (0.02 - 60 $s^{-1}$) and a stress range (0.5 to 100 mPa·s) and Newtonian viscosities were obtained up to a concentration of $C = 0.085$ %. Above this last value, the flow curves were not linear anymore. On Fig.1, we display the flow curves fitted with a linear curve for a shear rate increase and then, for a shear rate decrease. Viscosities were obtained from a linear fit $\sigma = \eta\dot\gamma$.

#### Cone-plane rheometry rheometry

For the cone-plane geometry, after an initial shear at 100 $s^{-1}$, the initially applied shear-rate is 10 $s^{-1}$. The

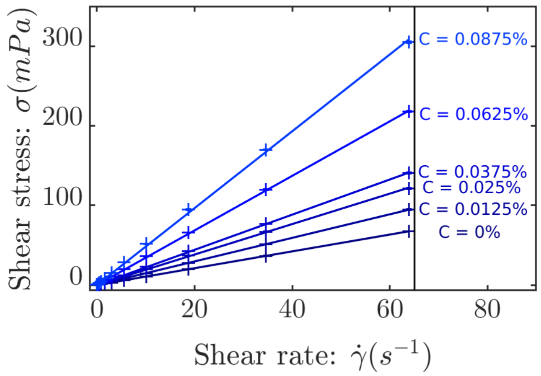


FIG. 1. Flow curves $\sigma(\dot\gamma)$, obtained from low-Shear rheometry. Data were fitted with a linear curve $\sigma = \eta\dot\gamma$ to extract the viscosity $\eta$.

rotation rate is then increased by steps via 1 min steady rotation rates $\omega_R$. The corresponding averaged shear-stresses are measured up to a shear rate of 100 $s^{-1}$. Then, the shear rates are successively decreased in reverse conditions, essentially to verify how measurements are reproducible. The corresponding flow curves are reported on Fig. 2B and Fig. 2C.

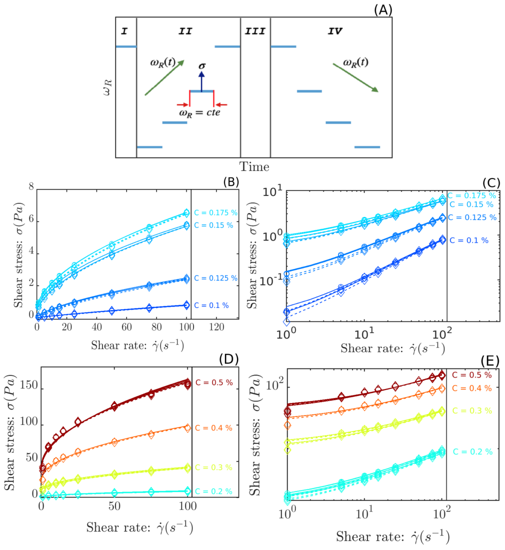


FIG. 2. Cone-plane rheometry. (A) Sketch of the shearing protocol. The flow curves $\sigma(\dot\gamma)$, are obtained through successive increases (diamonds) and decreases (circles) of the shear-rate values corresponding to changes in the rotation rates $\omega_R$. The data are fitted with the three parameter curve of equation 1. (B) Flow curves in the range 0.1 % $C < 0.175$ % on a linear scale, (C) corresponding in $log - log$ scale. (D) Flow curves in the range of 0.2 % $C < 0.5$ % on a linear scale and (E) in $log - log$ scale respectively.

#### Aging of the Carbopol mixture

Carbopol mixtures can present a small aging effect on a large time scale, as can be seen in Figure 3, where the shear stress $\sigma$ evolves as a function of time, presenting a minimal deviation after 30 min. However, it should be noted that the aging effect does not affect the performance of the characterization of the bacteria swimming in Carbopol, because all samples containing bacteria and Carbopol are changed every 25 min, where many times to characterize only one concentration, at least 4 fresh samples were used.

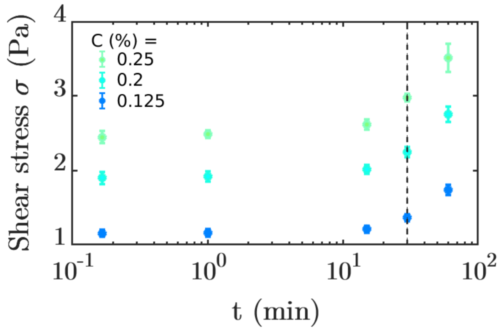


FIG. 3. Aging rheology. Shear stress as a function of time for Carbopol at concentrations C = 0.1 %, C = 0.2 % and C = 0.25 %. Black dashed line represent the maximum time to change a Carbopol mixture with bacteria added to perform 3D tracking experiments.

#### Non-linear flow curve fitting

In the moderate range of concentrations we operate, $0\,\% < C < 0.5\,\%$, we describe the rheology of the mixture with a fitting function bridging a Newtonian viscous behavior at low Carbopol concentrations, to the rheology at higher concentrations where an Herschel-Bulkey (HB) flow-curve is expected. In the spirit of soft glassy materials rheology, we use a form with an exponent $n = 1/2$ characteristic of microgels suspensions and analyze the experimental rheological curves with the following empirical fitting relation:

$$\sigma = \sigma_0 + \eta \frac{\dot{\gamma}}{1 + (\tau_0 \dot{\gamma})^{1/2}} \quad (1)$$

where the parameter $De = \tau_0 \dot{\gamma}$ can be viewed as a Deborah number involving an internal reorganisation time scale $\tau_0$. At low shear $De \ll 1$, a linear Bingham-fluid form is obtained (eventually with a Newtonian rheology for $\sigma_0 = 0$) and for $De \gg 1$, this expression recovers the HB form with $n = 1/2$. In this picture the parameter $\eta$ can still be qualitatively interpreted, at least from a dimensional perspective, as a viscosity (HB parameter $a = \eta/\sqrt{(\tau_0)}$). On Fig.4, the corresponding viscous parameter $\eta$ and internal time scale $\tau_0$ are displayed as a function of the concentration. The quality of the fitting procedure can be tested as we plot the rescaled shear stress $\Delta\tilde{\sigma} = \frac{(\sigma - \sigma_0)\tau_0}{\eta}$ as a function of Deborah number $De$ and compare the results with the dimensionless function:

$$\Delta\tilde{\sigma}(De) = \frac{De}{1 + De^{1/2}} \quad (2)$$

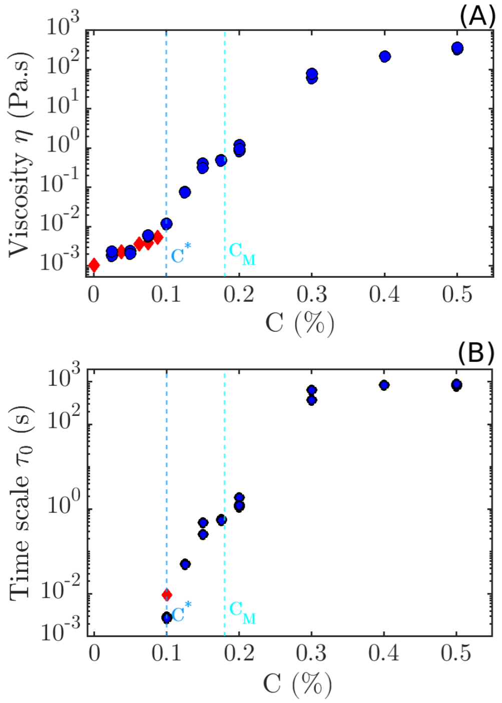


FIG. 4. Fitting parameters of the flow curves (see eq. (1)) as a function of the concentration, for $C > C^*$. Red diamonds are for the low-shear Couette rheometer and blue circles for the cone-plane rheometer. (top) viscous parameter $\eta(C)$ and (bottom) internal time scale $\tau_0(C)$. The yield stress values $\sigma_0(C)$ are represented in the main text.

### MOTILITY MEASUREMENTS EXTENDED DATA

#### Test for a diffusivity proxi in the ballistic-diffusive regime

In the finite time observation range we used, sometimes, MSD measurements essentially display a regime of pure ballistic motion. However, from the time correlation decay of the swimming orientation, a persistent swimming orientation time can be measured. If the stochastic process in homogeneous in time, the resulting random

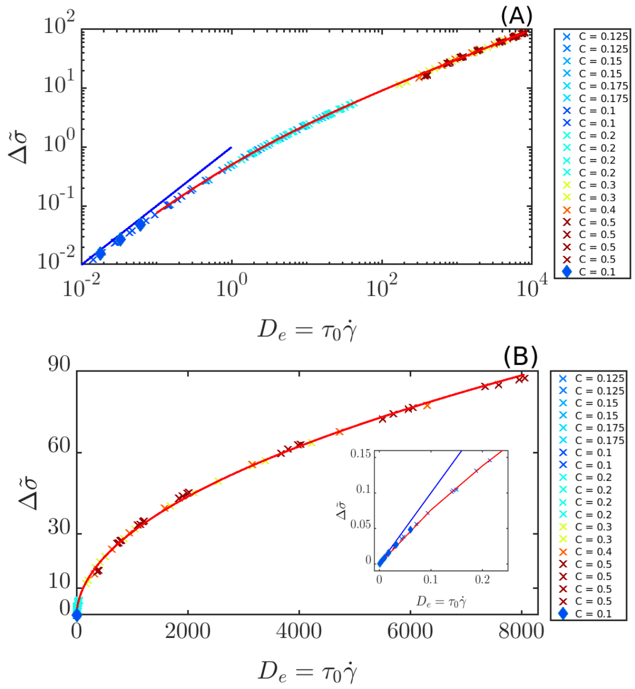


FIG. 5. Rescaled shear stress $\Delta\tilde{\sigma}$ as a function of Deborah number $De$ in logarithmic and linear scales. (A), Log-log scale representation and (B) same data in a linear representation. The inset corresponds to Deborah numbers in the cross-over region between the linear ($De$) and the HB ($De^{1/2}$ limiting regimes. Diamonds correspond to low-shear and crosses to the cone-plane data. Concentrations $C$ are labelled in percent (%). The red line is the function $F(De) = De/(1 + De^{1/2})$

walk will display a diffusivity $D$ with a value:

$$D = \frac{V_b^2 \tau_p}{3} \quad (3)$$

where $V_b$ is the bacterium velocity and $\tau_p$ the persistence time. This expression is chosen as a proxi for the bacterium diffusivity when the tracks appear as pure-ballistic. For completion, we test this proxi for tracks in the ballistic diffusive regime where from the MSD calculation a diffusivity could be extracted. This test $D_{BD}$ vs $\frac{V_B^2\tau_p}{3}$ is presented on Fig.3.

### OPTICAL DETECTION OF MICROSCOPIC HETEROGENEITY

To identify the presence of optical heterogeneity, droplets of Carbopol mixtures are confined between two glass slides separated by a height $H = 200\,\mu$m. Using white light at a moderate level of intensity the samples essentially transparent are visualized at 63X in the the low pixel intensity level (see Fig. 7(A)). We use as a reference image the pure motility buffer and subtract it to the images taken in the same conditions, at different Car-

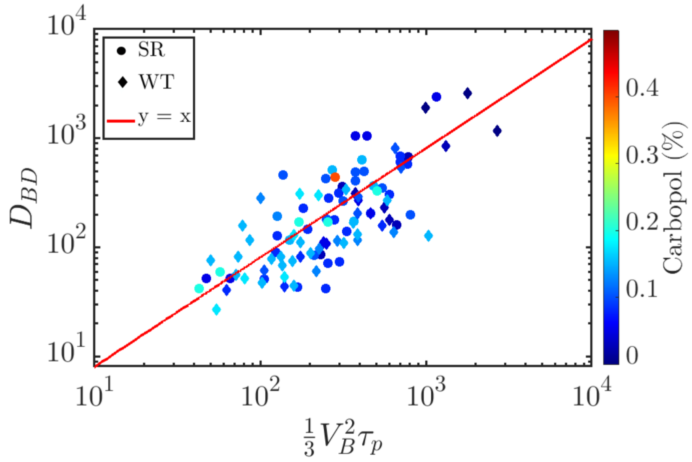


FIG. 6. Test of the proxy value of eq.(3) for wild-type and smooth-runner in the ballistic-diffusive regime : $D_{BD}$ as a function of $\frac{V_B^2\tau_p}{3}$.

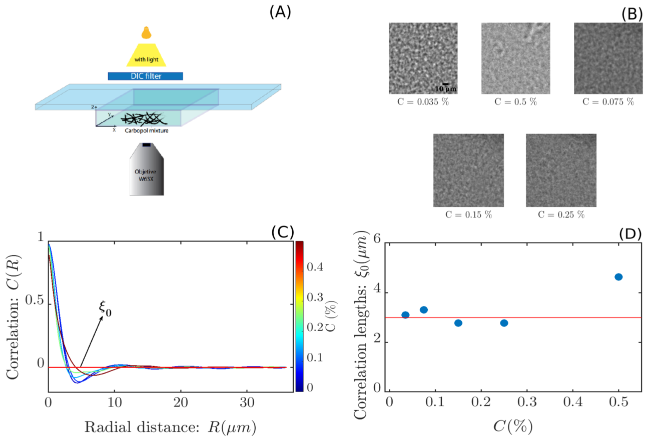


FIG. 7. Detection of optical heterogeneities. (A) Set-up schematics showing a C-Apochromat 63x/1.2W objective and a (DIC) filter illumination of the cabopol mixture. (B) resulting images after post processing (see text) displaying contrast heterogeneities at different Carbopol concentrations C. The scale bar is 6 $\mu$m. (C) Image intensity spatial correlations $C(R)$ as a function of radial distance $R$ at different Carbopol concentrations $C$. Correlation lengths parameter $\Lambda_0$ corresponding to the passage of the correlation function at zero as a function of Carbopol concentration. The straight line corresponds to the mean value for $C < 0.4\,\%$.

bopol concentrations. By adjusting the contrast in post-processing the presence of globular patterns is identified (Fig. 7 (B)). Quantitatively, the intensity spatial correlation $C(R)$ is computed at different concentrations (Fig. 7(C)). The correlation functions display typical damped oscillation features characteristic of the disordered globular structure which can be quantitatively characterized by the passage to zero at $\xi_0$ such as to define a structural scale $\lambda_0 = 2\xi_0$, which corresponds to half the distance between white and black extrema. We thus identify a typical diameter for these globular shape heterogeneities: $\langle\xi\rangle = 6\,\mu$m.

[1] G. Junot, T. Darnige, A. Lindner, V. A. Martinez, J. Arlt, A. Dawson, W. C. K. Poon, H. Auradou, and E. Clément, Phys. Rev. Lett. **128**, 248101 (2022).
[2] S. K. DeWitt and E. A. Adelberg, Genetics **47**, 577 (1962).
